# *Khdc3* Regulates Metabolism Across Generations in a DNA-Independent Manner

**DOI:** 10.1101/2024.02.27.582278

**Authors:** Liana Senaldi, Nora Hassan, Sean Cullen, Uthra Balaji, Natalie Trigg, Jinghua Gu, Hailey Finkelstein, Kathryn Phillips, Colin Conine, Matthew Smith-Raska

## Abstract

Genetic variants can alter the profile of heritable molecules such as small RNAs in sperm and oocytes, and in this manner ancestral genetic variants can have a significant effect on offspring phenotypes even if they are not inherited. Here we show that wild type female mice descended from ancestors with a mutation in the mammalian germ cell gene *Khdc3* have hepatic metabolic defects that persist over multiple generations. We find that genetically wild type females descended from *Khdc3* mutants have transcriptional dysregulation of critical hepatic metabolic genes, which persist over multiple generations and pass through both female and male lineages. This was associated with dysregulation of hepatically-metabolized molecules in the blood of these wild type mice with mutational ancestry. The oocytes of *Khdc3*-null females, as well as their wild type descendants, had dysregulation of multiple small RNAs, suggesting that these epigenetic changes in the gametes transmit the phenotype between generations. Furthermore, injection of serum from wild type mice with ancestral history of *Khdc3* mutation into wild type females is sufficient to cause hepatic transcriptional dysregulation in their offspring. Our results demonstrate that ancestral mutation in *Khdc3* can produce transgenerational inherited phenotypes, potentially indefinitely.

## Introduction

It is becoming increasingly apparent that non-DNA molecules inherited from germ cells contribute significantly to the transmission of traits and diseases across generations. This has been demonstrated in response to a variety of exposures such as diet or stress in model organisms (1, 2) and has been confirmed in human epidemiological studies (3, 4). The mechanisms driving this process remain poorly defined and are complicated by the erasure of acquired epigenetic molecules such as DNA methylation early in embryonic development. Expression of germ cell small RNAs such as microRNA (miRNA) and tRNA fragments (tsRNA) are responsive to exposures and inherited at fertilization, making them potential candidates for passage of phenotypic information across generations (1, 5). Microinjection of these small RNAs from the germ cells of an exposed mouse into an unexposed embryo is sufficient to recapitulate the descendant’s phenotypes, demonstrating causality (1). However, the absence of an RNA-dependent RNA polymerase in mammals suggests that small RNAs by themselves cannot propagate phenotypes across multiple generations.

Non-inherited ancestral genetic variants can also affect descendants’ risk of disease, as has been demonstrated in cancer (6, 7), thyroid hormone metabolism (8), body weight (9), anxiety (10), type I diabetes (11), and folate metabolism (12). This phenomenon is often referred to as “genetic nurture” based on the effect of non-inherited DNA variants nurturing phenotypes in the next generation (13). In all described cases, the observed phenotypes disappear after a few generations. The molecular mechanisms driving these cross-generational traits, in the absence of inheritance of the causal variant, remains largely undescribed, with the exception of paramutation effects that are driven by inherited RNA molecules (14, 15). Some of the genes associated with genetic nurture act by altering the expression of RNAs in the gamete, which can affect processes such as glucose metabolism and body weight (16). Other reported examples involve genes that do not have any known function in the germ cells, and likely alter descendants’ phenotypes indirectly, by changing underlying physiology in a manner that subsequently alters germ cell heritable molecules, not dissimilar from the mechanisms of exposure-based changes to heritable germ cell molecules (8).

These experiments are typically performed in males to avoid the confounding influence of in utero- and breastmilk-mediated effects. As a result, examination of oocyte small RNAs in the inheritance of phenotypes has been much less studied, because of the confounding influence of the placenta and breastmilk, and also because the dozens of oocytes per mouse yield much less RNA than can be obtained from the millions of sperm isolated from a male. However, in vitro fertilization has been utilized to demonstrate that oocytes are also capable of transmitting exposure-based non-genetic information across generations (17).

*Khdc3* is a mammalian gene expressed in the male and female germ cells that encodes a protein containing an RNA-binding KH domain that localizes to a multi-protein complex at the oocyte periphery known as the subcortical maternal complex (18, 19). *Khdc3*-null female mice have decreased fertility caused by defects in maintaining euploidy during embryogenesis (20). Human females with homozygous mutations in the ortholog *KHDC3L* are infertile, which has been associated with abnormal DNA methylation in oocytes (18). *KHDC3L* (and *Khdc3)* does not localize to the nucleus and does not contain a DNA methyltransferase domain, suggesting that the DNA methylation defects are secondary outcome and not the main function of this gene.

We demonstrate that wild type female descendants of *Khdc3*-null mice have dysfunctional expression of hepatic metabolic genes that persists over multiple generations, and that this effect is transmitted from both male and female mutant ancestors. This corresponds with abnormal levels of hepatic metabolites in the serum of these wild type mice with mutant ancestors. The persistence of these abnormalities in genetically wild type mice suggests that altered epigenetic information in the germ cells is transmitting the inherited metabolic phenotypes. Accordingly, we observed that the oocytes of *Khdc3*-null females and their wild type descendants have multiple dysregulated miRNAs and tsRNAs, suggesting a mechanism of inheritance.

## Results

### Wild type females descended from Khdc3-null ancestors display hepatic transcriptional dysregulation

Global transcriptome analysis of *Khdc3*-null oocytes revealed significant dysregulation of genes important in metabolic processes regulated by the liver, especially the metabolism of lipids (atherosclerosis, PPAR signaling, and pantothenate and CoA metabolism) and of carbohydrates (glycolysis/gluconeogenesis, and fructose and mannose metabolism) (Fig. 1A) (21–24). Despite the abnormal expression of metabolic genes in the *Khdc3*-null oocyte, this gene is only expressed in female reproductive tissue, with no detectable expression in the liver and minimal expression in other metabolic tissues such as adipose or pancreas (Fig. 1B).

**Figure 1.**
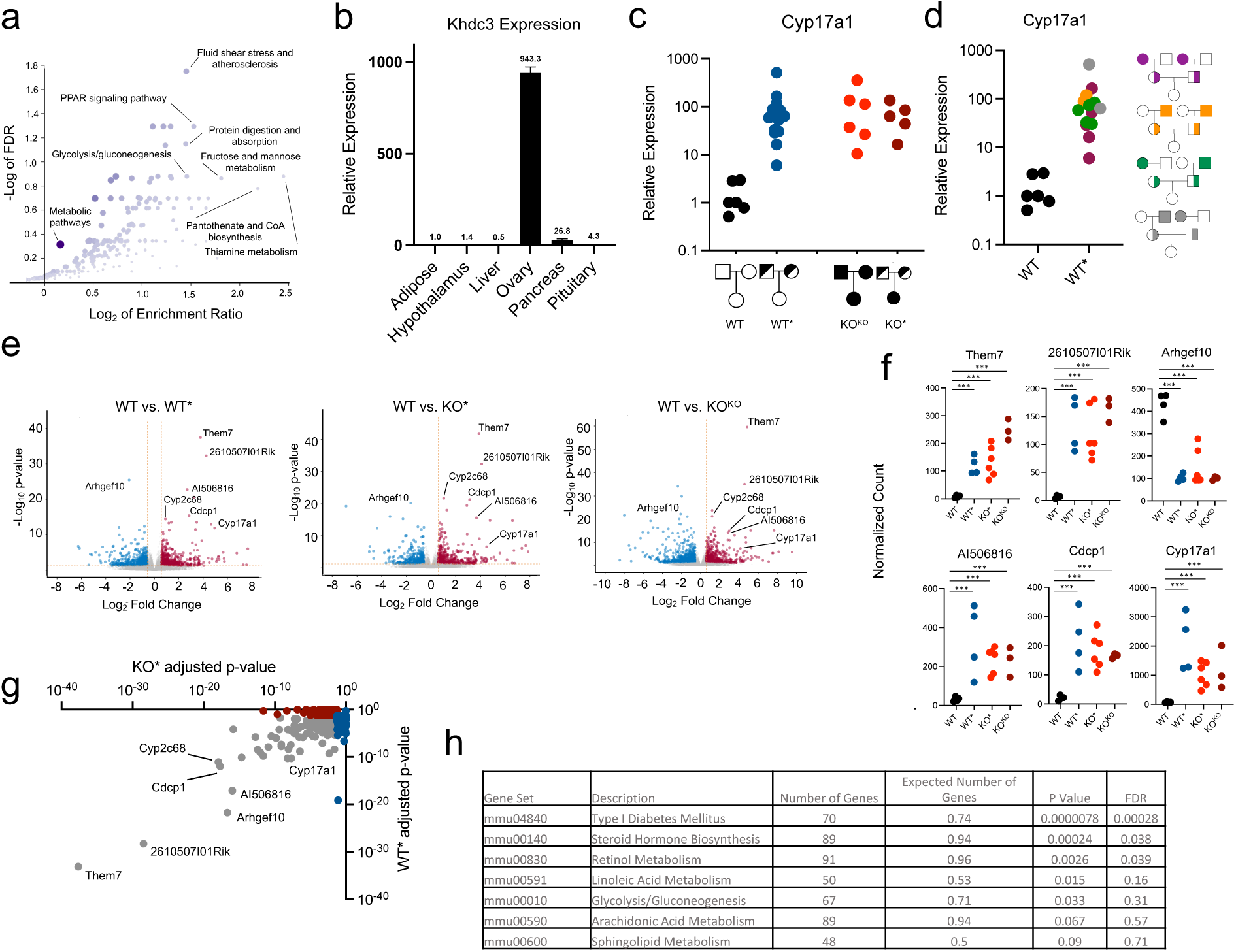
Female mice with an ancestral history of Khdc3 mutation manifest hepatic transcriptional dysregulation that is independent of genotype. (a) Gene ontology scatterplot for dysregulated genes in *Khdc3-*null oocytes, based on RNA-Seq. The y-axis represents −log FDR (false discovery rate) and the x-axis represents log_2_ enrichment ratio. Dysregulation of genes important in lipid and carbohydrate metabolism are highlighted. (b) Murine expression of *Khdc3* in various tissues, detected by qPCR (N=3). (c) Relative mRNA expression of *Cyp17a1* in the livers of WT, WT*, KO^KO^, and KO* mice measured by qPCR. Each dot represents an individual mouse. (d) Relative mRNA expression of *Cyp17a1* in the livers of WT* female mice generated from various male and female *Khdc3*-null grandparents measured by qPCR. (e) Volcano plots depicting differentially expressed genes identified by RNA-seq in the livers of WT vs. WT*, KO* and KO^KO^ mice, respectively (N=4-6). Red dots represent upregulated genes in WT*, KO* and KO^KO^ mice while blue dots represent genes that were downregulated in WT*, KO* and KO^KO^ mice. The x-axis shows log_2_ fold change values, and the y-axis denotes −log_10_ p-values. (f) Dot plots of common dysregulated liver genes amongst the WT*, KO*, and KO^KO^ mice compared to WT mice; ***p-adjusted < 1 × 10^−5^. (g) Dot plot of the significantly dysregulated genes in KO* livers (x-axis) versus WT* livers (y-axis), based on adjusted p-value. Red dots reveal significantly dysregulated genes in KO* mice, blue dots reveal significantly dysregulated genes in WT* mice, and grey dots represent commonly significantly dysregulated genes in both KO* and WT* mice. (h) Gene ontology of the most common dysregulated genes identified in the livers of both WT* and KO* revealed abnormalities in pathways critical for lipid and glucose metabolism.

Expression of *Cyp17a1*, a gene central to lipid metabolism (25, 26), was increased in the livers of *Khdc3*-null (knockout, or “KO”) females, despite a lack of expression or any known function of *Khdc3* in wild type livers. Elevated expression of *Cyp17a1* was observed in KO females generated from KO parents (referred to as “KO^KO^”) as well as KO females generated by mating *Khdc3* heterozygote parents (referred to as “KO*”) (Fig. 1C). *Cyp17a1* expression was also significantly increased, to the same extent as KO^KO^ and KO* females, in genetically wild type females descended from heterozygous parents (referred to as “WT*” to indicate genetically wild type mice that have descended from *Khdc3* heterozygous mutant parents and a combination of wild type and *Khdc3*-null grandparents). Thus, having an ancestor that carried a *Khdc3* mutation was associated with elevated *Cyp17a1* expression, and the *Khdc3* genotype had no effect on *Cyp17a1* expression, revealing a DNA-independent form of inheritance (Fig. 1C). Increased *Cyp17a1* mRNA expression was observed in WT* females generated from all possible combinations of male and female KO grandparents (Fig. 1D). In addition to conventional genotyping, the wild type genotype of these mice was confirmed with RNA-Seq of ovaries (Supplemental Figure 1).

RNA-Seq of female WT* livers revealed a pattern of global transcriptional dysregulation that overlapped significantly with KO* and KO^KO^ females (Fig. 1E-F), demonstrating that shared ancestry rather than genotype accounts for the transcriptional abnormalities in WT* mice. This is supported by the observation that most significantly dysregulated genes in WT* livers were similarly dysregulated in KO* livers (Fig. 1G). Gene ontology of the common dysregulated genes in livers of WT* and KO* mice revealed enrichment in metabolic processes in which the liver plays a central role, especially lipid and glucose metabolism (24–27) (Fig. 1H).

### WT* defects persist over multiple generations

When WT* male and female mice were mated with each other to generate the next generation (referred to as WT**, signifying the 2^nd^ generation of genetically wild type mice), hepatic transcriptional dysregulation of metabolic genes persisted. There was a similar pattern of dysregulation in WT** females derived from either male or female *Khdc3*-null ancestors (denoted WT**(P) and WT**(M), respectively, denoting <u>P</u>aternal or <u>M</u>aternal ancestral mutant history) (Fig. 2A-B). Comparison of dysregulated genes between WT**(P) and WT**(M) females showed significant overlap (Fig. 2C) revealing that the same pattern of abnormal gene expression is observed in wild type females descended from both male and female mutant ancestors.

**Figure 2.**
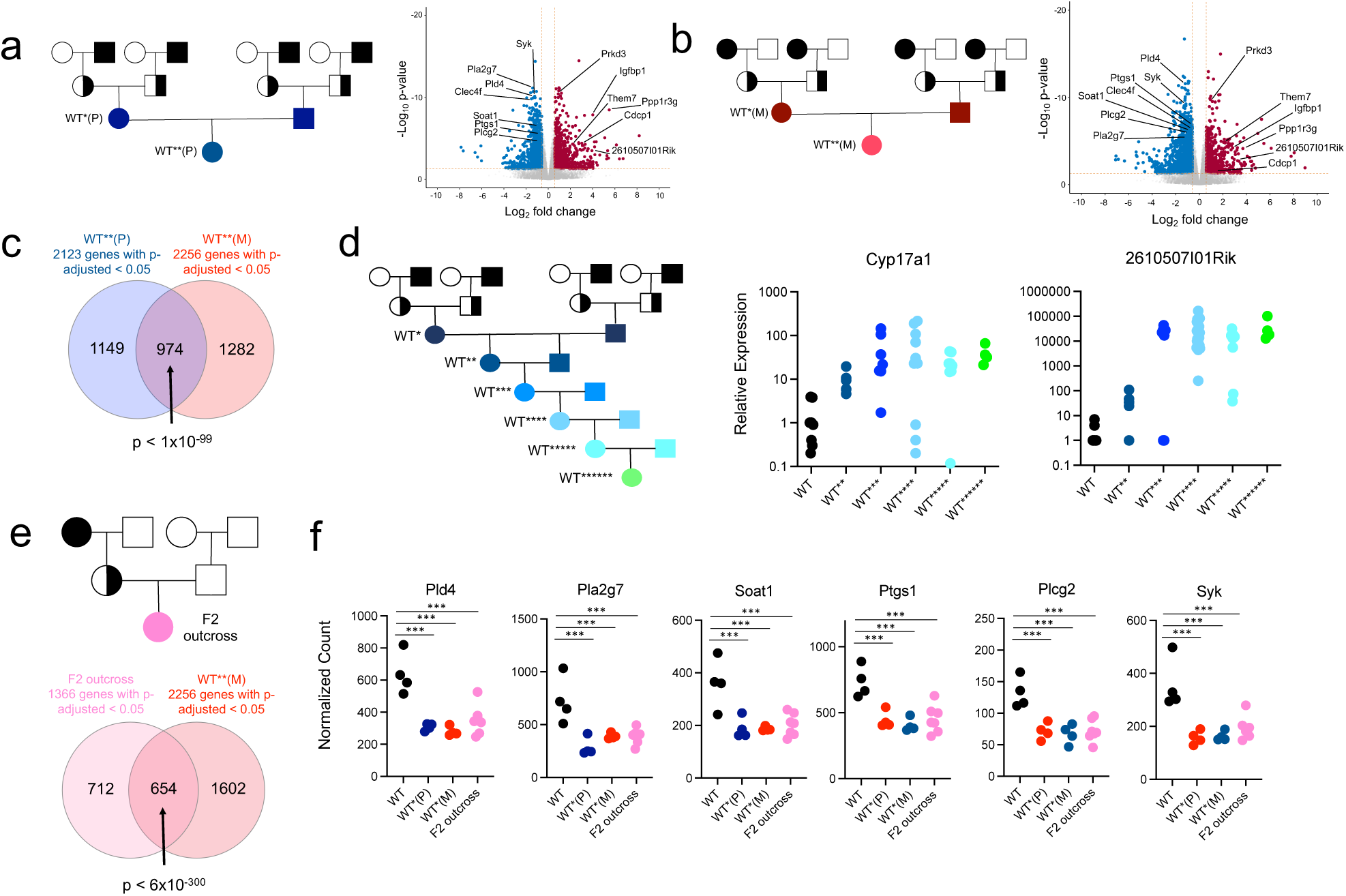
WT* defects persist over multiple generations and can be passed through the maternal and paternal ancestral lines. (a) Schematic of experimental mating to form WT**(P) mice. *Khdc3-*null mice are represented by black filled shapes, WT mice represented by white filled shapes, and *Khdc3*-heterozygote mice represented by half-filled black shapes. Volcano plot displaying dysregulated genes in the livers of WT**(P) mice compared to WT mice (N=4). Red dots represent upregulated genes and blue dots represent downregulated genes in the WT**(P) mice. (b) Schematic of experimental mating to form WT**(M) mice. Volcano plot displaying dysregulated genes in the livers of WT**(M) mice compared to WT mice (N=4). Red dots represent upregulated genes and blue dots represent downregulated genes in the WT**(M) mice. (c) Venn diagram representing the number of significantly dysregulated liver genes of WT**(P) and WT**(M) female mice compared to WT mice. (d) Pedigree schematic representing mating of WT*(P) male and female mice over successive generations to form WT******(P) mice. Dot plots represent relative liver mRNA expression of *Cyp17a1* and *2610507I01Rik* of the 2^nd^ through 7^th^ generation WT female mice descended from male *Khdc3*-null mice as measured by qPCR. (e) Pedigree schematic representing creation of a F2 outcross WT female from the mating of a female *Khdc3*-null female and WT male and subsequent F1 generation *Khdc3*-heterozygote female with WT male. Venn diagram depicting the overlap of commonly dysregulated genes in the livers of F2 outcross WT and WT**(M) female mice compared to WT mice (N=4-7). (f) Dot plots representing liver gene dysregulation amongst WT, WT*(P), WT*(M), and F2 outcross WT female mice. ***p < 1 × 10^−5^.

WT(P) male and female mice were mated with each other in order to examine passage these defects over successive generations. Descendants of male mutants were examined to avoid potential confounding from non-germ cell mechanisms of inheritance such as mitochondrial inheritance, fetal-maternal communication across the placenta, or transmission of molecules via breastmilk. Expression of *Cyp17a1* and *2610507I01Rik*, two genes dysregulated in WT*, KO*, KO^KO^, and WT**(P) female mice, demonstrated persistent and worsening dysregulated expression in the 2^nd^ through 6^th^ generation of females, in which each additional asterisk (*) denotes the successive number of genetically wild type generations (Fig. 2D). In a minority of mice, expression reverted back to WT levels (Fig. 2D). Importantly, those mice that reverted to wild type levels of *Cyp17a1* maintained elevated expression of *2610507I01Rik*, and vice-versa, suggesting that the abnormal expression of these genes is driven by independent units of inheritance.

*Khdc3*-null females were outcrossed with a true wild type male without any ancestral history of *Khdc3* mutation, in order to examine the persistence of the observed defects through only the maternal line without any contribution from males that descended from mutants. 100% of the F2 outcrossed females, descended from a maternal grandmother mutant, had persistent transcriptional dysregulation of metabolism-related genes in the liver (Fig. 2E). Many of these dysregulated genes overlapped with WT* females descended from both male (WT*(P)) and female (WT*(M)) homozygous-null ancestors (Fig 2F). In this mating scheme, persistence of hepatic transcriptional dysregulation in all 7 examined females is not consistent with mechanisms of inheritance involving in-cis transmission, including an unaccounted DNA polymorphism/mutation, DNA methylation, or histone protein modification, in which any chromosome from the *Khdc3*-null grandmother would have been inherited by ∼50% of the F2 outcross generation.

### Metabolic phenotype of WT* females

Litter sizes from WT* male and female matings were not significantly different from litter sizes of true wild type matings (Fig. 3A). Furthermore, there was no difference in body weight at birth, 3 weeks, and 8 weeks of age between WT** and WT females (Fig. 3B), revealing that the detected hepatic transcriptome changes are not the result of altered litter sizes or growth rates. Metabolic phenotype analysis in 8 month-old females revealed no significant differences between WT**** and WT mice in body weight, fat composition, food intake, or energy expenditure (Fig. 3C).

**Figure 3.**
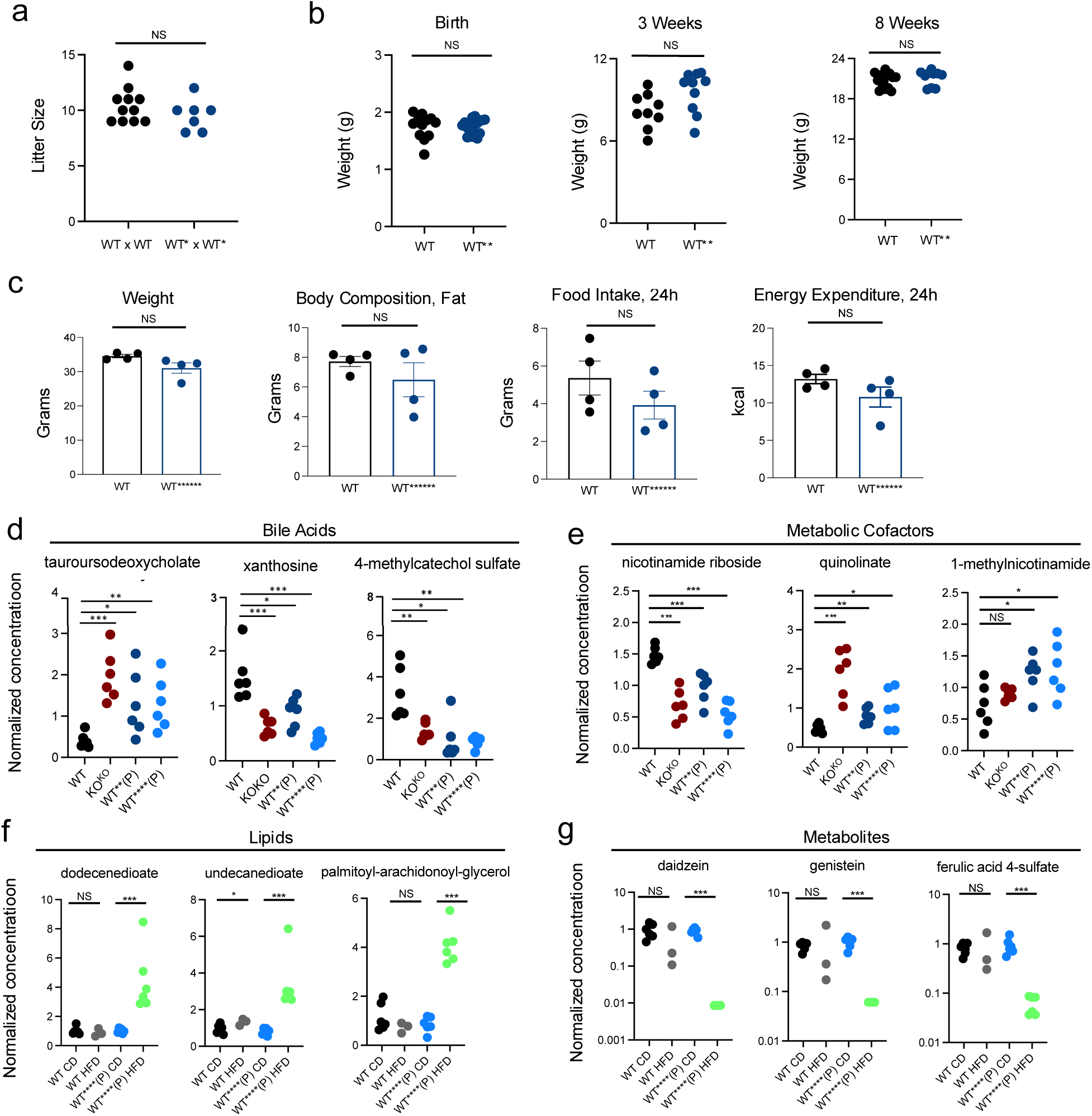
Wild type mice with ancestral history of *Khdc3* mutation have dysregulation of multiple hepatically-metabolized molecule in the serum. (a) Litter sizes in wild type (WT x WT) and WT* (WT* x WT*) matings. (b) Weights of WT** pups at birth, 3 weeks, and 8 weeks. (c) Metabolic phenotype of WT****** females at 8 months of age, including weight, fat composition, food intake, and energy expenditure. (d-e) Dot plots revealing concentration of bile acids and metabolic cofactors in KO^KO^, WT**(P), and WT****(P) female mice compared to WT mice. (f) Dot plots revealing metabolic cofactors in WT****(P) female mice exposed to a HFD compared to WT**** mice fed a conventional diet and WT mice fed a HFD. (g) Dot plots revealing metabolites in WT****(P) female mice exposed to a HFD compared to WT**** mice fed a conventional diet and WT mice fed a HFD. *p < 0.05, **p < 0.005, ***p < 0.0005.

Global serum metabolomic analysis was performed to examine the metabolic consequences of the observed transcriptional dysregulation. Both WT** and WT**** female mice had abnormal levels of multiple bile acids, which are synthesized in the liver (28–31) (Fig. 3D). The magnitude of dysregulation was similar to that observed in KO^KO^ females (Fig. 3D), suggesting that mutational ancestry is more important than the organism’s genotype. These females also had abnormal concentrations of various metabolic cofactors (Fig. 3E) that are associated with hepatic metabolism (Fig. 3E) (32–35).

WT**** mice were metabolically challenged with 8 weeks of a high fat diet (HFD). When compared with true wild type mice exposed to a HFD, WT**** females exposed to a HFD showed a significant increase in multiple lipid molecules that were not dysregulated in WT**** females that consumed a conventional diet, or in WT mice consuming a HFD (Fig. 3F). Thus, there were latent metabolic abnormalities that could not be detected unless stressed with a HFD. The HFD WT**** serum also had decreased levels of multiple other hepatic metabolites (36–38) some of which were also decreased in WT HFD females although at a much lesser magnitude (Fig. 3G). These findings demonstrate that the hepatic transcriptional dysregulation observed in these mice has significant effects on multiple important metabolites found in the blood.

### Small RNA dysregulation

Livers of WT** females had dysfunctional expression of multiple small RNA-processing genes, especially genes that regulate tRNA (*Trmt9b* (*39*), *Ang* (*40*), *Nsun6* (*41*), *Elac1* (*42*)) (Fig. 3A) and miRNA processing (*Mettl1* (*43, 44*), *Ago2* (*45*), *Exosc10*, *Syncrip* (*46*)) (Fig. 4A). Based on this observation, in combination with the fact that gamete small RNAs can transmit non-genetically inherited phenotypes in mice, we hypothesized that the observed phenotypes were driven by the defective inheritance of small RNAs from the germ cells of *Khdc3* mutant ancestors. Small RNA-Seq of KO^KO^ oocytes revealed dysregulation of multiple miRNAs and tRNA fragments, with minimal piRNA or rRNA dysregulation (Fig. 4B). In WT**(P) oocytes, there was also abnormal expression of tRNA fragments and miRNA, most of which were different small RNAs than the ones dysregulated in KO^KO^ oocytes (Fig. 4C). Of note, the WT**(P) oocytes had downregulation of the tRNA fragment Gly-GCC, for which expression in sperm has been associated with inherited metabolic and hepatic gluconeogenesis phenotypes in offspring (1, 47, 48). There were 3 tRNA fragments and 18 miRNAs that were commonly dysregulated in both KO^KO^ and WT**(P) oocytes (Fig. 3D), suggesting that their abnormal expression is established in KO^KO^ oocytes and is not normalized with reintroduction of a wild type *Khdc3* allele. The function of most of these commonly dysregulated tsRNAs and miRNAs remains undescribed, however miR-107, which was upregulated in both KO^KO^ and WT**(P) oocytes, is involved in hepatic lipid metabolism (49).

**Figure 4.**
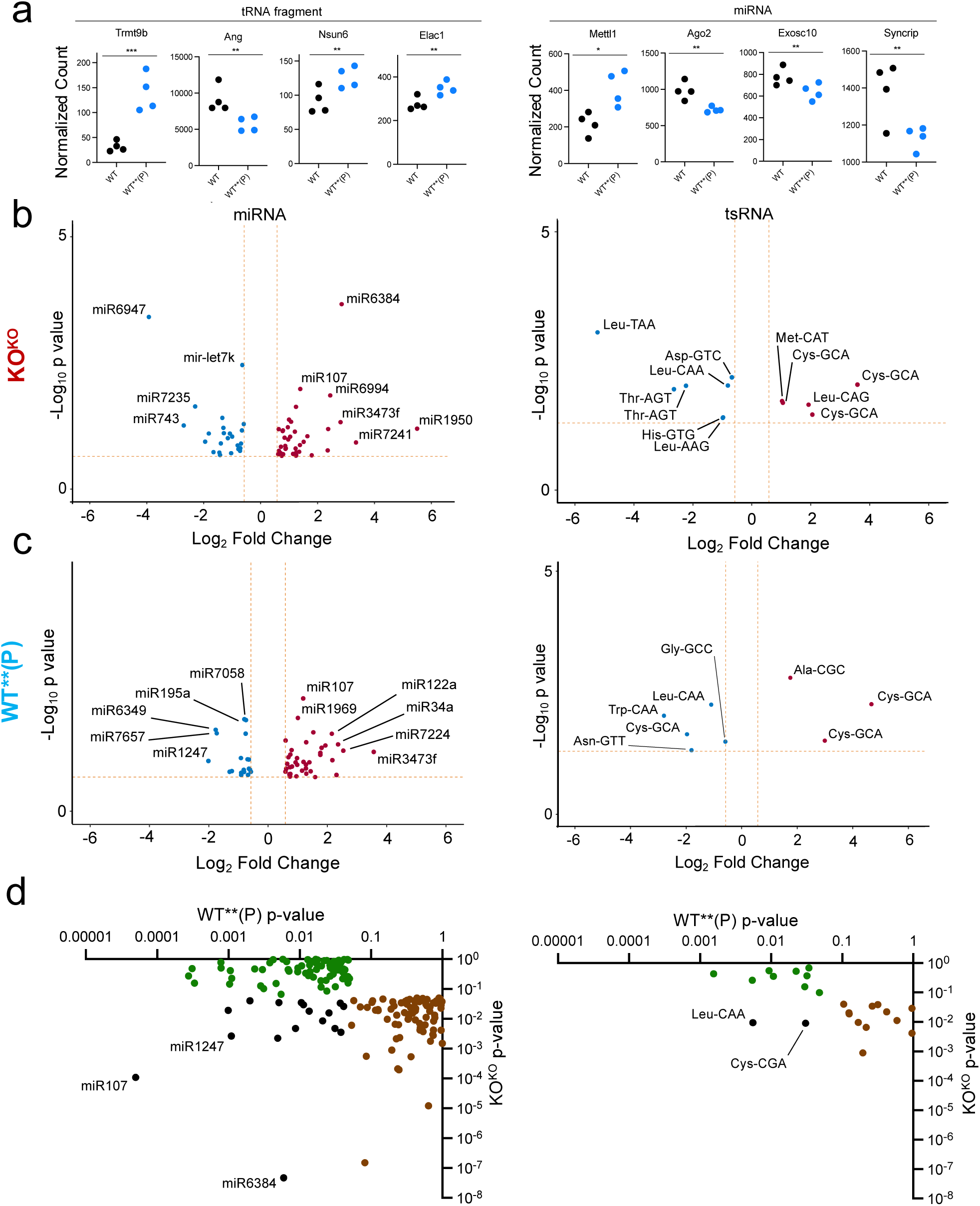
Oocytes of KOKO and WT**(P) mice have dysregulated expression of multiple miRNAs and tsRNAs. **(a)** Dot plots representing dysfunctional expression of small RNA-processing genes of the livers of WT**(P) mice as measured in RNA-seq**. (b)** Volcano plots depicting differentially expressed miRNAs and tRNA fragments in KO^KO^ oocytes compared to WT oocytes. Red dots represent upregulated small RNAs, and blue dots represent downregulated small RNAs (N=4). **(c)** Volcano plots depicting differentially expressed miRNAs and tRNA fragments in WT**(P) oocytes compared to WT oocytes. Red dots represent upregulated small RNAs, and blue dots represent downregulated small RNAs (N=4). **(d)** Scatterplot of the most significantly dysregulated small RNAs of WT**(P) oocytes (x-axis) versus KO^KO^ oocytes (y-axis), based on p-value. Green dots reveal dysregulated miRNAs and tRNA fragments in WT**(P) oocytes, brown dots reveal dysregulated miRNAs and tRNA fragments in KO^KO^ oocytes, and black dots represent commonly dysregulated small RNAs in both WT**(P) and KO^KO^ oocytes.

### Serum from Wild Type Mice with Ancestral History of Khdc3 Mutation is Sufficient to Alter Hepatic Gene Expression in Offspring of Wild Type Mice

Recent studies have demonstrated that circulating factors in the blood can induce intergenerational phenotypes that recapitulate the effects observed in an exposure-based model (50). When serum from WT* mice (genetically wild type with Khdc3-null ancestry) was injected intraperitoneally into wild type females that were subsequently mated, the offspring manifested an altered hepatic transcriptome (Fig. 5A-B). This observation suggests that yet-unidentified factors carried in the serum are sufficient to drive the observed cross-generational changes. Of note, some genes were commonly dysregulated in both this serum transfer experiment and the WT* female mice, for example Cyp17a1. However, there were other genes that did not overlap between the two experiments, likely because the serum transfer approach cannot reproduce the duration and concentration of exposure to serum-based factors that occurs in wild type descendants of *Khdc3* mutants.

**Figure 5.**
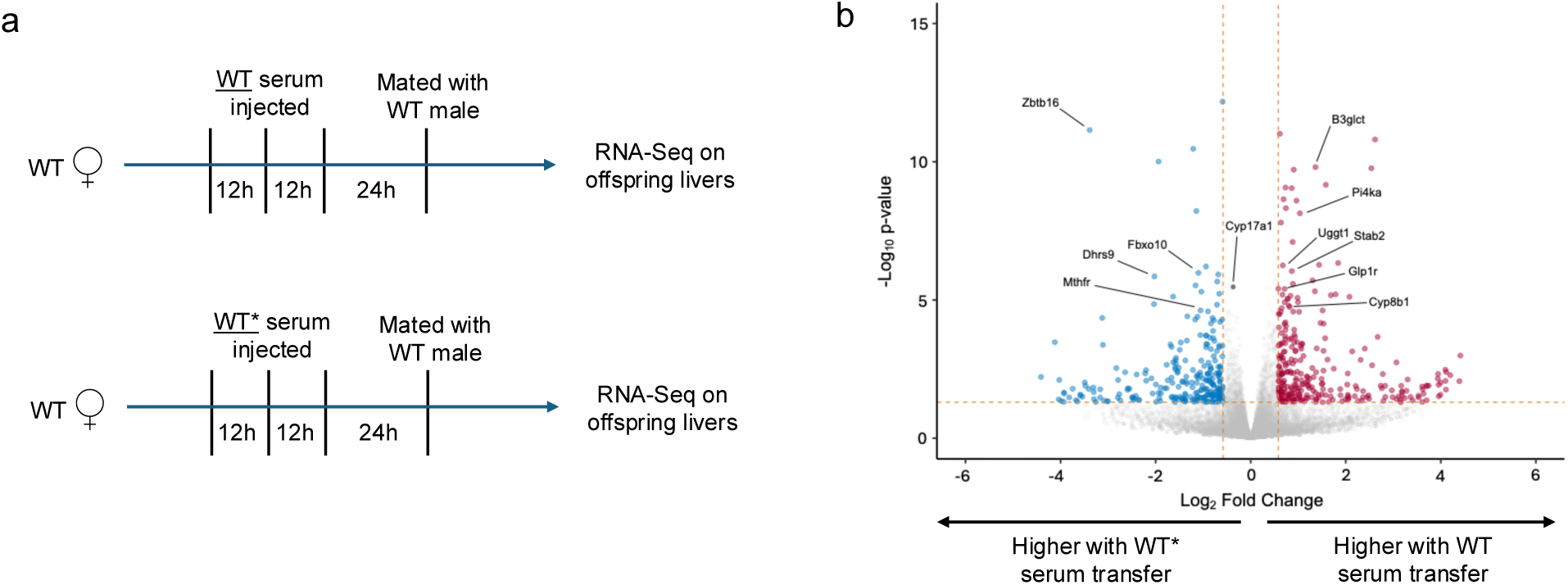
Injection of serum from KO mice in WT females is sufficient to cause transcriptional dysregulation in WT offspring. (a) Schematic of serum transfer experiment. (d) Volcano plot depicting differentially expressed genes in the wild type offspring born to mothers who received injection with either WT or KO serum. *p < 0.05, **p < 0.005, ***p < 0.0005.

## Discussion

The presence of abnormal phenotypes in wild type organisms descended from mutant-carrying ancestors has been described in mice and humans. This phenomenon can occur from mutations that affect epigenetic phenomenon in the germ cell (6, 7, 51), but in other cases the mutation affects a process that has no clear connection with heritable molecules in the germ cell, such as with thyroid metabolism or type I diabetes (8, 11). In all previously described examples, the observed defects disappear after one or two generations.

We find that loss-of-function mutation in *Khdc3* alters the hepatic metabolism of female wild type descendants over at least 6 generations in a manner that cannot be rescued with re-introduction of the wild type *Khdc3* allele. This is paralleled by abnormal levels of multiple metabolites in the serum of these mice. Because there is no detectable *Khdc3* expression in the liver, we suspect that the defects in WT* mice are caused by inherited molecules that affect hepatic metabolism independent of the inheritance of a functional *Khdc3* gene. We expect that other metabolic tissues such as the pancreas and adipose will also demonstrate evidence of metabolic dysregulation.

We utilized an outcrossing scheme to demonstrate that the observed defects persist when passed only through the maternal line. In this mating scheme, 100% of the observed F2 outcrossed female mice had evidence of hepatic metabolic dysregulation. This pattern of inheritance is not consistent with in-cis mechanisms such as an unaccounted DNA variant, DNA methylation, or histone protein modification, because in these scenarios the alleles in the *Khdc3*-null maternal grandmother would be present in ∼50% of the F2 outcrossed females.

We chose to examine females because of the reported fertility phenotypes in *KHDC3* human (18, 19) and *Khdc3* mouse female mutants (20). Human *KHDC3L*-null females are infertile, while mouse *Khdc3*-null females are subfertile. The fertility defects are associated with oocyte DNA methylation abnormalities that we suspect are sequelae of a more primary defect in mutant organisms. There are no reports of abnormal phenotypes in *Khdc3*-null males, however our detected metabolic abnormalities could affect males as well, which will be an important future investigation. Furthermore, it will be important to examine how *Khdc3*-null mice transmit exposure-based information across generations to affect phenotypes in non-exposed offspring, as has been observed in numerous studies, most thoroughly with a HFD or stress.

We have demonstrated that *Khdc3*-null oocytes have abnormal expression of multiple miRNAs and tsRNAs, some of which persist over multiple generations in WT* mice, providing a potential mechanism of inheritance. Microinjection experiments using sperm RNA have demonstrated that small RNAs are sufficient to drive metabolic derangements in offspring (1, 52), however the maintenance of defects to the successive generation remains unexplained. The amount of RNA needed for these experiments currently prevents using oocyte RNA in such an experimental paradigm, because of the low total RNA yield obtained from the oocytes of a single mouse.

A recent study demonstrated that transfer of serum from a stress-exposed mouse into a non-exposed mouse can recapitulate the effects observed in the exposed’s offspring, suggesting serum factors can modify the composition of heritable molecules in the germ cells and affect offspring traits (50). Indeed, we observed that the serum from a WT* mouse is sufficient to cause hepatic dysregulation when it was injected into a wild type female/mother. The observation that serum-based factors can drive cross-generational hepatic transcriptome changes suggests that the metabolic alterations in each generation with an ancestral history of *Khdc3* mutation drive changes to heritable molecules in the oocyte, which then drive metabolic dysregulation in the next generation.

One important conclusion from studies on genetic nurture is that wild type mice descended from mutant ancestors should be used thoughtfully and with caution, because phenotypes can be obscured that would be apparent with the use of wild type mice without any ancestral history of mutation. In sum, the effect of non-inherited genetic variants on offspring phenotypes remains a relatively unexplored and potentially significant contributor to traits and disease risk.

## Materials and Methods

### Mice

FVB/N WT (Jax 001800) mice were purchased from The Jackson Laboratory and bred in-house. Frozen *Khdc3-null* sperm was obtained from the Mutant Mouse Resource and Research Center (MMRRC, North Carolina) and the *Khdc*3 KO mouse was rederived. Female mice were used at 8 weeks of age. In individual experiments, all animals were age-matched. All mice were maintained under specific pathogen-free (SPF) conditions on a 12-hour light/dark cycle, and provided food and water ad libitum. All mouse experiments were approved by, and performed in accordance with, the Institutional Animal Care and Use Committee guidelines at Weill Cornell Medicine.

### Mouse Genotyping

As described in this study, WT mice were generated from WT parents. WT* and KO* mice were generated from the mating of *Khdc3* heterozygote parents that were themselves generated from mating of WT and KO^KO^ mice. KO^KO^ mice were generated from *Khdc3* KO parents. DNA was extracted from tail biopsies by incubating the tails biopsies in an alkaline lysis buffer (24mM NaOH and 0.2mM disodium EDTA) for 30 minutes at 95°C and then on ice for 10 minutes. A neutralization buffer (40 mM Tris-HCl) was added to the samples and diluted 1:10 with water. PCR to detect *Khdc3* deletion was performed with 50–200 ng of DNA using the Phire Green Hot Start II DNA Polymerase Master Mix (Thermo Scientific) in the Bio-Rad T100 Thermal Cycler. PCR for *Khdc3* deletion was performed using a forward primer for the wild type allele F1, a separate forward primer that incorporates the deletion allele F2, and a common reverse primer R1. The primers were as follows: P1: 5’-TGCCTGGGCAGGTTATTTAG-3’, P2: 5’-CGAGCGTCTGAAACCTCTTC-3’ and P3: 5’-AGCTAGCTTGGCTGGACGTA-3’. P1 and P2 amplified the wild type allele and P1 and P3 amplified the *Khdc3* mutant allele (KO). The PCR products were separated on a 1% agarose gel with SYBR safe DNA gel stain. Because of the different sizes of the PCR products, the genotypes can be easily determined from the band patterns on DNA gels.

### Liver RNA Extraction

Total RNA extraction was performed on the liver of 8 week old mice using the Qiagen RNeasy Lipid Tissue Mini Kit (Qiagen) according to the manufacturer’s instructions. Samples were eluted in 30μl nuclease-free water. Nucleic acid concentration, A260/280 and A260/230 ratios was determined via NanodropD-1000 (Thermo Scientific). Extracted samples were aliquoted and stored at -80°C.

### RT-PCR and quantitative RT-PCR

2000ng RNA per sample was used to generate cDNA using SuperScript III First-Strand Synthesis System (Invitrogen) following the manufacturer’s instructions. cDNA was diluted to 1:4 in water before performing qPCR using SYBR Green PCR Master Mix (Applied Biosystems). qPCR reactions were performed on a QuantStudio 6 Flex Real Time PCR Instrument. Cycling conditions were as follows: Initial denaturation 95°C for 3 minutes, 40 cycles of denaturation at 95°C for 15 seconds followed by annealing/extension at 60°C for 60 seconds. Relative expression was calculated using the ΔΔCt method, using YWHAZ as the reference gene. Primers used are listed in Supplemental Table 1.

### Oocyte RNA Isolation

The oocytes were dissected following a published protocol with some modifications (53). Briefly, the ovaries were first dissected from unstimulated 8-week-old WT and *Khdc3*-null mice and placed in PBS. The ovary was dissected from surrounding para-ovarian fat and subsequently placed in a 35-mm culture dish with 2mL PBS and 20μl collagenase and 20μl DNase in a 37°C incubator for 20 minutes. Using a dissecting microscope, the oocytes were separated from granulosa cells, theca cells, and stromal cells, and transferred to a petri dish with PBS using a mouth-controlled micropipette. RNA was extracted from the oocytes using the PicoPure RNA Isolation kit (Thermo Fisher) following the manufacturer’s instructions and eluted in 20 µL of the provided elution buffer.

### Liver RNA-seq Library Preparation

Sequencing libraries were prepared using the Illumina TruSeq Stranded Total RNA kit according to the manufacturer’s protocol and sequenced to a depth of 40 million reads per sample. The paired-end (PE) libraries were sequenced on Novaseq platform.

### Oocyte RNA-seq Library Preparation

Sequencing libraries were prepared using the SMART-Seq v4 Ultra Low Input RNA plus Nextera XT DNA Sample Preparation according to the manufacturer’s protocol and sequenced to a depth of 40 million reads per sample. The paired-end (PE) libraries were sequenced on Novaseq platform.

### RNA-seq Bioinformatics

All sequenced reads were assessed for quality using FastQC (54). Adapter trimming and filtering of low-quality bases (<20) were performed using cutadapt (55). The trimmed reads were then aligned with Hisat2 (56) against the mouse reference (GRCm38) genome. Per gene counts were computed using htseq-count (57) (genocode M15). DESeq2 (58) bioconductor package was used for normalization and differential gene expression analysis.

### Oocyte small RNA isolation

Eight week old female mice were sacrificed and 30 oocytes were dissected under stereomicroscopy from the ovaries of each mouse, as described above. Four separate WT, *Khdc3*-null, and WT(P)** RNA samples were generated from separate mice. Total RNA was isolated using a magnet-based method (ChargeSwitch Total RNA Cell Kit).

### Oocyte small RNA-Seq Library Preparation

Small RNA quality and concentration was assessed using an Agilent Bioanalyzer and small RNA Chip to ensure a clear small RNA population of more than 1ng. Subsequently, small RNA libraries were generated and sequenced to a depth of 10 million reads per sample. Small RNA libraries were generated using the NEXTFLEX Small RNA-seq Kit, which were sequenced on the Novaseq platform.

### Small RNA-seq Bioinformatics

Read quality was assessed using FastQC (v0.11.8), and adapter sequences were trimmed using Trimmomatic (v0.39). After adapter trimming, reads were mapped sequentially to rRNA mapping reads, miRbase, murine tRNAs, pachytene piRNA clusters (59), repeatmasker and Refseq using Bowtie 2 alignment algorithm (v2.3.5) and totaled using Feature Counts on the Via Foundry (v1.6.4) platform (60). To assess for differentially expressed small RNAs , data was loaded into R Statistical Software and analyzed using the DESeq2 package (58). Differentially abundant small RNAs were determined as those with a log fold change > 0.58 and P-value < 0.05.

### Serum preparation

Blood was collected from the submandibular (facial) vein of mice. Approximately 400-500μl of whole blood was collected from each mouse and placed in an SST amber microtainer blood collection tube (BD). Blood collection tubes were placed in a 37°C incubator for 30 minutes, followed by 10 minutes at 4°C. Samples were then centrifuged at 5500 RPM for 10 minutes at 4°C. The serum was collected from the top of the tube and stored at -80°C until needed.

### Metabolic Phenotpying

Body weight, body fat composition, food intake, and energy expenditure of 8 month old female mice were measured using the Promethion Core System.

### Metabolomic profiling

Metabolomics profiling was conducted using ultra-high-performance liquid chromatography-tandem mass-spectrometry by the metabolomics provider Metabolon Inc. (Morrisville, USA) on mouse serum samples from WT, WT**(P), WT****(P) and KO^KO^ mice, as well as these mice after consuming a high fat diet. The metabolomic dataset measured by Metabolon includes known metabolites containing the following broad categories – amino-acids, peptides, carbohydrates, energy intermediates, lipids, nucleotides, cofactors and vitamins, and xenobiotics. Statistical differences were determined using unpaired t-test.

### High fat Diet

Female WT and WT****(P) mice were divided into two diet groups, one group receiving a high-fat diet (HFD, D12492i; Research Diets Inc., New Brunswick, NJ) and the other group received a normal diet with 10% fat (D12450Ji; Research Diets Inc., New Brunswick, NJ). All mice were given access to food and water *ad libitum* and were maintained on a 12:12-h light-dark artificial lighting cycle. After 8 weeks of each diet, serum was collected and stored in the -80°C.

### Serum Metabolites

Serum lipids were quantified using Beckman Coulter kits OSR6516, OSR6296, and OSR6295. Serum proteins were precipitated using trichloroacetic acid. The pellet was then dissolved, tryptic digested, desalted, and analyzed by LC-MS/MS for protein identification. The data were processed by MaxQuant. MS data was searched against Uniprot mouse protein database.

### Serum Transfer

Serum was obtained from WT and WT* (WT females descended from Khdc3-null ancestors), as described above. Pooled WT serum was injected intraperitoneally into 3-month old female WT (n=2) and WT* (n=2) mice. Similarly, pooled WT* serum was injected IP into female WT (n=2) and WT* (n=2) mice. IP injections consisted of 200μl pooled serum every 12 hours for 3 doses (each mouse received a total of 600μl injected serum). The female WT and WT* mice were subsequently mated with a WT male 24 hours later. Livers from female mice of each offspring were dissected at 1 month of life. RNA was extracted and RNA sequencing was performed on the samples, as described above.

### Statistical Analysis and data visualization

Gene ontology performed with WebGestalt (https://www.webgestalt.org/) using KEGG analysis. Volcano plots generated with ggVolcanoR (https://ggvolcanor.erc.monash.edu). Venn diagrams generated with InteractiVenn (http://www.interactivenn.net/). Scatter plots were generated with Prism 9.2.0. The manuscript was written using Microsoft Word v16.66.1. Figures were assembled on Microsoft PowerPoint v16.66.1

## Data Availability

All RNA-Seq and small RNA-Seq data are available in the NCBI GEO database, accession numbers GSE281631 (oocyte RNA-Seq, Figure 1), GSE 282184 (liver RNA-Seq, Fig. 1), GSE283285 (liver RNA-Seq, Fig. 2), GSE288429 (F2 outcross liver RNA-Seq, Figure 2), GSE 287350 (oocyte small RNA-Seq, Figure 4), and GSE288414 (serum transfer RNA-Seq, Figure 5).

## Acknowledgements

This work was supported by the Young Investigator Award (LS) at Weill Cornell Medicine, Pediatric Scientist Development Program under the National Institute of Child Health under award number K12HD000850 (SC), Friedman Family Clinical Scholar Award (MSR), and National Institute of Child Health and Human Development (NICHD) of the National Institutes of Health under award number P50HD104454 (MSR).

## Author Contributions

L.S. and M.S.R. conceived and planned the experiments. L.S., S.C., N.H., H.F., K.P., and M.S.R carried out the experiments. U.B., N.T., J.G., and C.C. performed bioinformatic analysis of the RNA-Seq and small RNA-Seq data. L.S. and M.S.R. prepared the manuscript. All authors provided critical feedback and contributed to shaping the research, analysis, and manuscript.

## Additional Information

Supplementary Information is available for this paper.

## Supplemental Material

**Supplemental Figure 1.**
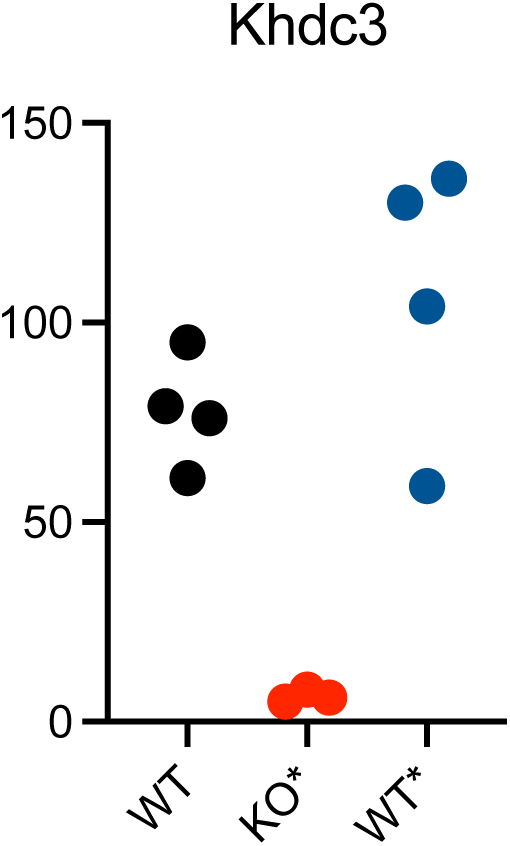
K*h*dc3 expression in ovaries of WT, KO*, and WT* mice, from RNA-Seq.

**Supplemental Table 1:**
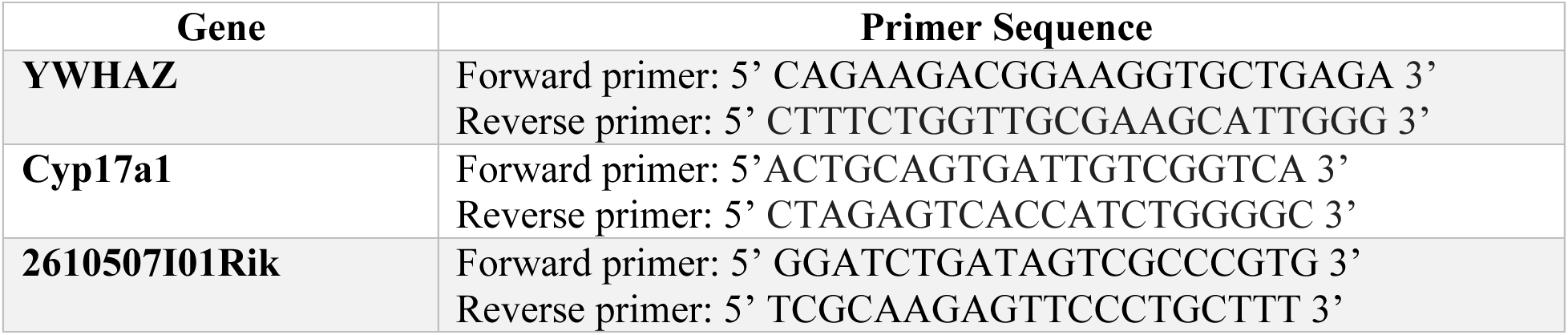
Primer sequences used for qPCR.

